# Sex-specific effects of social bonds on glucocorticoids in wild chimpanzees

**DOI:** 10.64898/2026.02.03.703594

**Authors:** Ellen D. Dyer, Zarin P Machanda, Megan F. Cole, Stephanie A. Fox, Maggy Kobusingye, Matthew Lem, Emily Otali, Richard W. Wrangham, Martin N. Muller, Melissa Emery Thompson

## Abstract

Across social species, strong social bonds are linked to health benefits, including reduced disease risk and increased survival. Chronic glucocorticoid exposure, frequently associated with negative health outcomes, may mediate this relationship, as bonds may reduce stressor exposure or buffer physiological responses. Among primates, these relationships have mostly been studied in female-bonded species, where kinship shapes cooperation and access to resources. Whether bonds confer simsilar benefits in species with different social structures – especially bonds between non-kin – remains unclear. We tested whether strong affiliative bonds were associated with glucocorticoid production in wild adult chimpanzees using 22 years of behavioural and urinary cortisol data. Bonds were quantified for same-sex adult dyads using an index of party association, proximity, and grooming. Bond strength was measured by averaging each individual’s top three bonds per year. Stronger bonds predicted lower cortisol in females but higher cortisol in males, a pattern that persisted after accounting for dominance rank, aggression, and contextual stressors. In this male-bonded species, stronger male bonds predicted higher physiological costs, while stronger female bonds, though weaker, appeared to attenuate stress. Our results challenge the assumption that social bonds are universally health-promoting, and suggest their physiological consequences vary with sex and social organization.

## Introduction

Social support is integral to human health and has been consistently linked to psychological and physical benefits [1–5].There is compelling evidence that social bonds promote cardiovascular health, impact cancer risk, and psychological well-being [5,6]. Lack of social support increases mortality risk as much as smoking 15 cigarettes a day and is even deadlier than obesity, physical inactivity, or exposure to air pollution [7]. Despite this, the mechanisms through which social bonds affect health remain unclear. In particular, we lack understanding of how biological pathways (*e.g*., stress physiology, inflammation) and social dynamics jointly shape health at various points in the life course [4,8].

The benefits of social bonds are not exclusive to humans. Studies of other social mammals demonstrate diverse fitness benefits from strong social ties, including improved reproductive success, enhanced offspring survival, and reduced mortality [9–11]. Recent studies from wild non-human primates have provided more specific evidence that social bonds impact mechanisms of aging and disease [2,11–13]. These model systems provide significant potential to disentangle the complex relationships between social environments and health because primate social relationships can be directly observed and objectively quantified across long time periods in a way that is not generally feasible for humans.

Social environments play an important role in exposure to stressors, thus the direct and indirect effects of these stressors are plausible routes by which social bonds could impact health. In response to a stressor, the hypothalamic-pituitary-adrenal (HPA) axis produces glucocorticoids which activate adaptive changes throughout the body to allocate energy and heighten arousal to allow the organism to respond to an immediate threat [4,14]. Because stress responses increase inflammation and draw resources away from maintenance processes, chronic stress and prolonged exposure to glucocorticoids are linked to an array of negative health outcomes [4,14,15]. Consequently, glucocorticoid exposure is proposed to be an important mechanism mediating the effects of social environments on health [14,16]. Additionally, the generalized nature of the stress response means that measuring glucocorticoids is a versatile tool for assessing the proximate biological effects of social contexts, which is essential for breaking down the long-term correlations with health outcomes [17].

Social bonds may reduce glucocorticoid exposure either by preventing exposure to stressors altogether or by mitigating maladaptive stress responses, helping individuals return to homeostasis following a challenge [3,8,18]. The latter mechanism—described in the literature as the *stress-buffering hypothesis*—is thought to influence physiology primarily during periods of acute stress [18]. While stress buffering is best documented in humans, where bond partners reliably dampen psychologically induced stress responses [5,18,19], there is also growing, albeit limited, evidence for analogous effects in nonhuman primates. Studies in baboons (*Papio spp.*), macaques (*Macaca spp.*), and chimpanzees (*Pan troglodytes*) show that close partners attenuate glucocorticoid production following naturally occurring social stressors, including the threat of infanticide [20], sudden social isolation [21], conspecific aggression [22], and intergroup encounters [23]. Social bonds may also reduce glucocorticoid exposure by altering the frequency or severity of stressors themselves. Strong partners can protect against harassment, reduce predation risk, and improve access to resources [8,24]—mechanisms widely supported across nonhuman primates that operate through direct social or ecological benefits.

In wild mammals, social bonds are most frequently linked to fitness outcomes related to reproduction, such as birth rates and infant survival [10,20,21,25]. But there is also evidence, primarily from wild non-human primates, that natural variation in social integration explains meaningful variation in HPA axis activity. Individuals with strong or focused social networks tend to exhibit lower glucocorticoid levels during periods of heightened stress, including environmental instability and social conflict [22,26–29]. Close partners can also help maintain physiological stability during daily challenges, particularly when their presence provides predictable social support [23,26]. Together, these results complement the stress-buffering patterns described above while also implying that the protective effects of social relationships extend beyond immediate reactions to acute threats.

Mechanistic insight comes from laboratory mammals, where social isolation produces robust neuroendocrine, behavioural, and immune dysregulation across taxa (reviewed in [30]). Isolation elevates glucocorticoids or alters ACTH responsiveness, causing acute cortisol spikes [30] as well as more persistent dysregulation [30–33]. Social deprivation can also impair immune function, slow wound healing or increase inflammation [30,31]. Captive primates show a similar overall pattern, though the direction of effects varies with developmental timing. Isolation can disrupt affiliative behaviour (*e.g*., marmosets, *Callithrix spp. [30,34]*) and HPA function: acute or repeated separation elevates cortisol (*e.g*., yellow baboons, *Papio cynocephalus*; juvenile rhesus macaques, *Macaca mulatta;* reviewed in [30]), whereas early-life isolation can lead to diminished basal cortisol years later (*e.g*., squirrel monkeys*, Saimiri sciureus;* rhesus macaques; [30]). Short-term individual housing of juvenile squirrel monkeys produces the opposite effect, with higher morning cortisol and heightened adrenal sensitivity to ACTH [30]. These findings demonstrate that while the specific endocrine outcomes vary, social isolation consistently disrupts HPA regulation across mammalian taxa.

Even during periods of relative stability, glucocorticoid production can covary with the intensity and stability of social bonds [29]. Researchers hypothesize that bonds provide predictability and a degree of control for individuals within social interactions, independent of any specific event [29,35]. However, the effects of social bonds may be entangled with other powerful influences on glucocorticoids, including social rank [35,36] and early life adversity [37,38]. A few reports in juvenile primates show contexts where investing in social bonds can increase glucocorticoids [39,40]. Juvenile blue monkeys (*Cercopithecus mitis*) and rhesus macaques with higher grooming rates or more peer interaction sometimes exhibit elevated glucocorticoids, suggesting that navigating uncertain or challenging relationships can be physiologically costly during development. However, among juveniles with more closely related peers, the number of strong ties and frequency of play bouts appeared to be linked to lower glucocorticoids [39,40], highlighting how the costs of social engagement shift with context.

Because these studies are overwhelmingly conducted over short time frames, it is, as yet, difficult to evaluate whether these proximate effects on stress responses are significant factors in the longer-term relationship between social bonds and health or survival. However, in a recent longitudinal report on wild female yellow baboons, high cumulative glucocorticoid exposure powerfully predicted age-specific mortality [41]. In the same study population, social bonds had previously been linked to both higher survival [42] and lower glucocorticoids [37], highlighting the importance of affiliative relationships. Like many cercopithecine monkeys, yellow baboons are female-philopatric, remaining in their natal groups throughout life and forming strong, matrilineal bonds that support cooperation in resource defense.

Research linking social bonds with health and survival has largely focused on these female-bonded cercopithecines, where social stability supports the formation and maintenance of long-term, kin-based affiliative relationships. By contrast, chimpanzees, which share a closer evolutionary relationship with humans, are male-philopatric and characterized by strong male-male bonds [43–45]. These bonds facilitate cooperation in agonistic contests over dominance status and mating opportunities [45,46] and can be leveraged to obtain mating concessions from high-ranking males [47–49]. Male bonds are also important for larger-scale cooperative networks used in hunting and territorial defense [50,51]. In contrast to female-philopatric cercopithecines, male-philopatric chimpanzees frequently form intersexual bonds with non-kin [52–54]. Female chimpanzees are less gregarious and form fewer and weaker bonds on average compared to either male chimpanzees or to female cercopithecines [44,52,53,55–58]. While females do groom and associate with males, these interactions are generally less stable and less affiliative than the long-term male-female ‘friendships’ documented in some cercopithecine species [59]. This might suggest an absence of fitness advantages that would otherwise promote bonding in female chimpanzees, although recent evidence demonstrates that females do reap the advantages of feeding tolerance and coalition formation from forming tolerant relationships with other females [60].

Here, we test the hypothesis that <u>strong bonds reduce glucocorticoid production</u> among wild Eastern chimpanzees (*Pan troglodytes schweinfurthii*). We assessed this with 22 years of longitudinal data on social relationships and urinary cortisol from the Kanyawara chimpanzee community in Kibale National Park, Uganda. Though fully habituated to the presence of researchers, the Kanyawara chimpanzees are not provisioned and receive medical intervention only in rare cases related to human impact (*e.g.*, snare removal). As such, they encounter a range of natural socioecological stressors upon which social bonds could act to impact fitness. This, in combination with their close evolutionary relationship to humans and complex social networks, makes wild chimpanzees a particularly relevant model system that can help address the current limitations in social support theory. We predicted that [**P1] chimpanzees with stronger affiliative bonds would have lower urinary cortisol concentrations than weakly-bonded chimpanzees of the same sex.** Since a number of sociodemographic and contextual variables (*e.g.*, age, dominance status, reproductive effort, social instability) are known to strongly impact cortisol levels in these chimpanzees [36,61–63] and may also be influenced by social bonds, we controlled for these variables to determine the independent effect of bond strength. Additionally, if the positive effects of social bonds result from increased resistance to stressors, we predicted that **[P2] under conditions where cortisol levels are typically elevated, strong social bonds would be associated with stronger cortisol-reducing effects**. We tested for interaction effects between social bonds and **(a)** unstable hierarchies; **(b)** mating contexts (swelling days for females, exposure to swollen parous females for males); and **(c)** high rank, a driver of elevated glucocorticoids for males. Chimpanzees, like humans, produce higher cortisol as they age [63]. In humans, social support is particularly critical in moderating HPA axis dysregulation with aging [64], thus we predicted that **[P3] the cortisol-reducing effect of social bonds would be stronger later in life.**

To test for one mechanism by which social bonds could influence cortisol, we considered whether individuals with strong social bonds experience fewer stressful events, as suggested in humans [3]. We focused on aggression as a potential mediator: if social bonds buffer stress by limiting exposure to aggression, then variation in aggression should account for some of the relationship between bond strength and cortisol. We predicted that **[P4] aggression mediates the effect of social bonds on cortisol.** In our prior work, aggression received had significant effects on female cortisol [62], while both aggression given and aggression received were important predictors for males [36].

## Methods

We analyzed variation in urinary glucocorticoid concentrations in adult chimpanzees (aged 15 years and older) in the Kanyawara community of wild, unprovisioned chimpanzees in Kibale National Park, Uganda, between Dec 1997 and Dec 2019. The Kanyawara community has been continuously monitored since 1987, with long-term demographic, behavioural, and ecological data collected following established protocols [65]. There were 9-13 males and 13-18 females in any given study year. An expanded methods section, including detailed field protocols, behavioural definitions, and hormone assay procedures, is provided in the electronic supplementary material to allow for brevity in the main text.

Immunoreactive cortisol was determined in urine samples via enzyme immunoassay with reagents and protocols (antibody R4866, University of California, Davis Clinical Endocrinology Laboratory) previously validated for this population [63]. Prior to analysis, all cortisol values were standardized for specific gravity and model-corrected for non-linear influences of time of day, year of assay, and sample storage time [36,63]. Pregnant females were excluded due to altered adrenal function during pregnancy [66].

Strength of social bonds and other behavioural measures were derived from 89,913 hours of observation, including group scans of party subgroup composition, all occurrences of agonistic behaviour, and an average of 829 hours of focal observation per individual (range: 5-2823 hrs) for 23 males and 28 females. Ordinal dominance ranks, standardized for cohort size (0 = low, 1 = high), were determined by daily Elo ratings [67] for males across the entire study and for females beginning in 2004. Sparser data on female agonistic interactions prior to 2004 required using Modified David’s scores [62,63].

To characterize the bondedness of adult males, we calculated annual dyadic composite sociality indices (DSI), following methods used elsewhere in primates [68]. The measure averages three component behavioural indices (party association, proximity, grooming) that are each standardized for the mean rate of interactions across all dyads of the same sex composition in the same year (average DSI = 1; see electronic supplementary material for full details on behavioural data collection, observation protocols, and calculation of component indices). *Bond strength* for males was defined as the average of the highest 3 DSIs (strongest 3 dyads) for each male in each year.

The DSI measure was not suitable for females because a large proportion of dyads failed to meet the minimum threshold of proximity (1 hour) necessary to compute the grooming index, and grooming was rare. Indeed, 84% of the annual female-female dyads in our study did not groom at all, and only 3% of female-female dyads groomed more than 3 times in a year. We substituted a dyadic association index (DAI), which was identical to the measure for males but without the grooming component. Thus, *bond strength* for females was defined as the average of the highest 3 DAIs for each year. Separately, we included a covariate for *grooming* which indicated how many of these top 3 social partners the female groomed with during the year (range 0-3). Minimum thresholds for data inclusion are described in the Supplement.

We performed linear mixed models in *R* (*lme4* and *car* packages) using each sample as the unit of analysis. To identify the independent effects of bond strength, we specified a model that included covariates identified as significant predictors of cortisol in prior studies of this population [36,63]. These included age, dominance rank, hierarchy instability, and mating context, log-transformed or centered, as appropriate. Hierarchy instability was defined categorically at the group level as stable or unstable. Following Muller et al. (2021)[36], we classified each day as stable or unstable based on whether any reversals in the relative ranks of two or more males occurred in the previous four weeks. Mating context was defined categorically by reproductive state for females, where females with maximally tumescent anogenital swellings were compared to cycling females without swellings and those in early or late lactation. For males, we defined mating context using a binary variable (0/1) as whether at least one parous, maximally-swollen female was observed in a party with the sampled male on the day of sampling. Where appropriate, fixed effects were log transformed (aggression) or centered on a meaningful zero value (Age = 15, Rank = 0.5, Bond strength = 1). Models additionally include random intercepts for chimpanzee ID and month-year of sample collection. We considered random slopes for rank and age, but due to convergence or singularity issues, neither was retained in the female models and only the slope for rank was included in the male models. Interactions between bond strength and other predictors were considered but only retained when they significantly improved the model fit, as determined with log-likelihood ratio tests. Multicollinearity was checked using *vif*, where all variance inflation factors were < 2. We additionally used the *DHARMa* package to evaluate goodness of fit. As commonly observed in large samples, these diagnostics identified a larger than expected proportion of outliers, though less than 1% and concentrated at low values. Despite this, the models were neither over- or under-dispersed nor zero-inflated and residuals were uncorrelated to predicted values.

Given that close kin are often among the strongest social partners in primates [69–71], we evaluated whether such relationships could be driving our results. In this study, no close kin dyads were observed among males, and among females, only mother–daughter dyads were present; no sister dyads were observed. Because these mother–daughter relationships were rare (one dyad in 1998 and one dyad from 2013–2016), we did not include kinship as a covariate in the main models. To ensure that these rare kin relationships were not influencing our results, we conducted a robustness check in which we removed only the individual-years in which a female was present with her mother or daughter (see electronic supplementary material for data). This approach preserves most of the data while eliminating potential confounding due to kinship.

After performing our primary analysis to detect the effects of bond strength on cortisol, we conducted a secondary analysis to examine whether these effects could be explained by variation in aggression. Specifically, we tested whether the apparent influence of bond strength was mediated by *aggression received* (rate per hour) for females, and both aggression received and *given* (rate per hour) for males, measured over the 14 days prior to and including the date of sampling. We selected a 14-day window to capture recent social interactions likely to influence current cortisol levels while avoiding excessively short-term fluctuations. This approach tests the idea outlined in the introduction: if strong social bonds buffer stress by reducing exposure to aggression, then including aggression in the models should reduce the observed effect of bond strength. These analyses were based on a smaller sample collected between 2005-2019, so we evaluated the mediation by comparing effect sizes to a similarly reduced version of the original model (reported in the supplement). The final dataset included 8003 complete data points for 28 adult females (representing 215 female-years) and 9560 data points for 21 adult males (211 male-years). The aggression-focused secondary analysis for aggression was based on 6730 samples for 23 adult females (177 female-years) and 7086 samples from 17 adult males (140 male-years).

## Results

### Strong bonds were associated with lower cortisol in females but higher cortisol in males (P1)

Increased bond strength, assessed via DAI, negatively predicted cortisol in female chimpanzees (estimate = -0.094, P = 0.0025, Table 1). Grooming did not explain additional variance in cortisol that was not already explained by association. Bond strength had the opposite effect on males; cortisol was positively predicted by DSI (estimate = 0.31, *P* = <0.001, Table 2). Despite some notable interactions with covariates, as below, the positive association between bond strength and cortisol persisted across all contexts for males. These results were robust to the presence of close kin: excluding female-years with a mother or daughter did not alter the main effects of bond strength or any interactions involving bond strength on cortisol (electronic supplementary material for detailed kinship robustness analyses).

**Table 1:** Female predictors of urinary’ glucocorticoid concentration. The table shows parameter estimates from the primary male model of urinary glucocorticoid concentrations for the full dataset of adult females. Bolded values indicate predictors with a statistically significant effect (p < 0.05).

|  | Primary Female Model |  |  |  |  |  |
| --- | --- | --- | --- | --- | --- | --- |
| *Model Predictors* | *Estimates* | *Variance* | *SE* | *SD* | *t* | *P* |
| Intercept | -0.007 |  | 0.062 |  | -0.11 | 0.913 |
| Age | **0.012** |  | **0.003** |  | **4.07** | **<0.001** |
| Rank | **0.256** |  | **0.117** |  | **2.19** | **0.030** |
| Instability | 0.059 |  | 0.031 |  | 1.90 | 0.058 |
| Reproductive State: Maximal Swelling | **0.238** |  | **0.050** |  | **4.80** | **<0.001** |
| Reproductive State: Early Lactation | -0.029 |  | 0.034 |  | -0.85 | 0.395 |
| Reproductive State: Late Lactation | -0.069 |  | 0.040 |  | -1.73 | 0.084 |
| Bond strength (DAI) | **-0.094** |  | **0.031** |  | **-3.00** | **0.002** |
| DAI x Rank | **-0.277** |  | **0.089** |  | **-3.11** | **0.002** |
| *Random Effects* |  |  |  |  |  |  |
| Chimp ID |  | 0.022 |  | 0.149 |  |  |
| Month/Year of sample collection |  | 0.176 |  | 0.420 |  |  |

N = 8003 urine samples, 215 female-years, 28 adult females
Random-effects variance components > 0 are interpreted as meaningful contributions of the grouping factor.

**Table 2:**
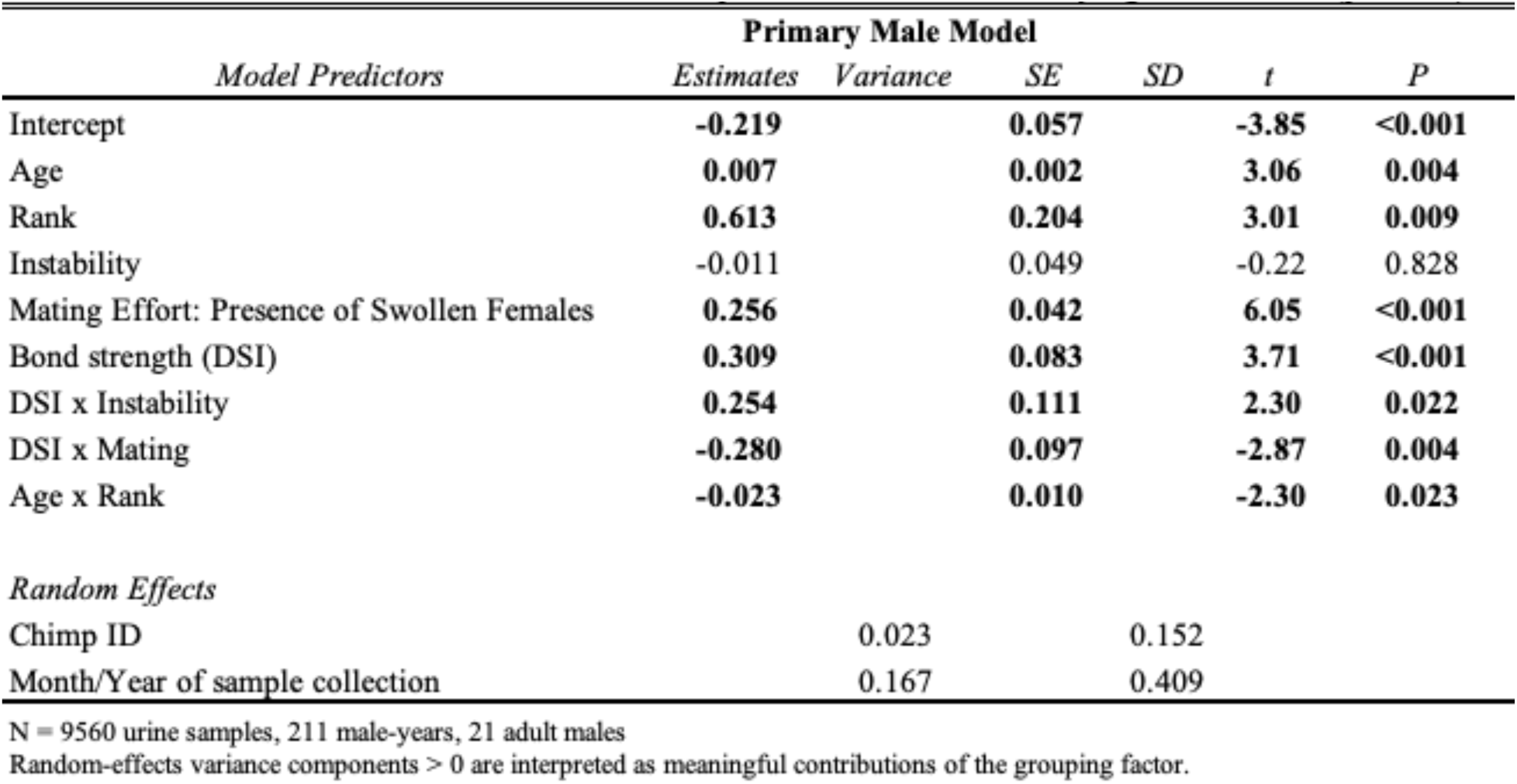
Male predictors of urinary glucocorticoid concentration. The table shows parameter estimates from the primary male model of urinary glucocorticoid concentrations for the full dataset of adult males. Bolded values indicate predictors with a statistically significant effect (p < 0,05).

### Strong bonds did not alleviate cortisol responses to stressful contexts (P2)

If bonds reduced vulnerability to stressors, we predicted stronger negative associations between bond strength and cortisol in contexts that reliably elicit stress responses. For females, neither hierarchy instability nor reproductive status moderated the negative effect of DAI on cortisol (Table 1). However, bond strength predicted a stronger reduction in cortisol for high-ranking vs. low-ranking females (Figure 1, estimate = -0.28, P = 0.002). Specifically, the highest cortisol levels were recovered from females who were high-ranking but weakly-bonded.

**Figure 1.**
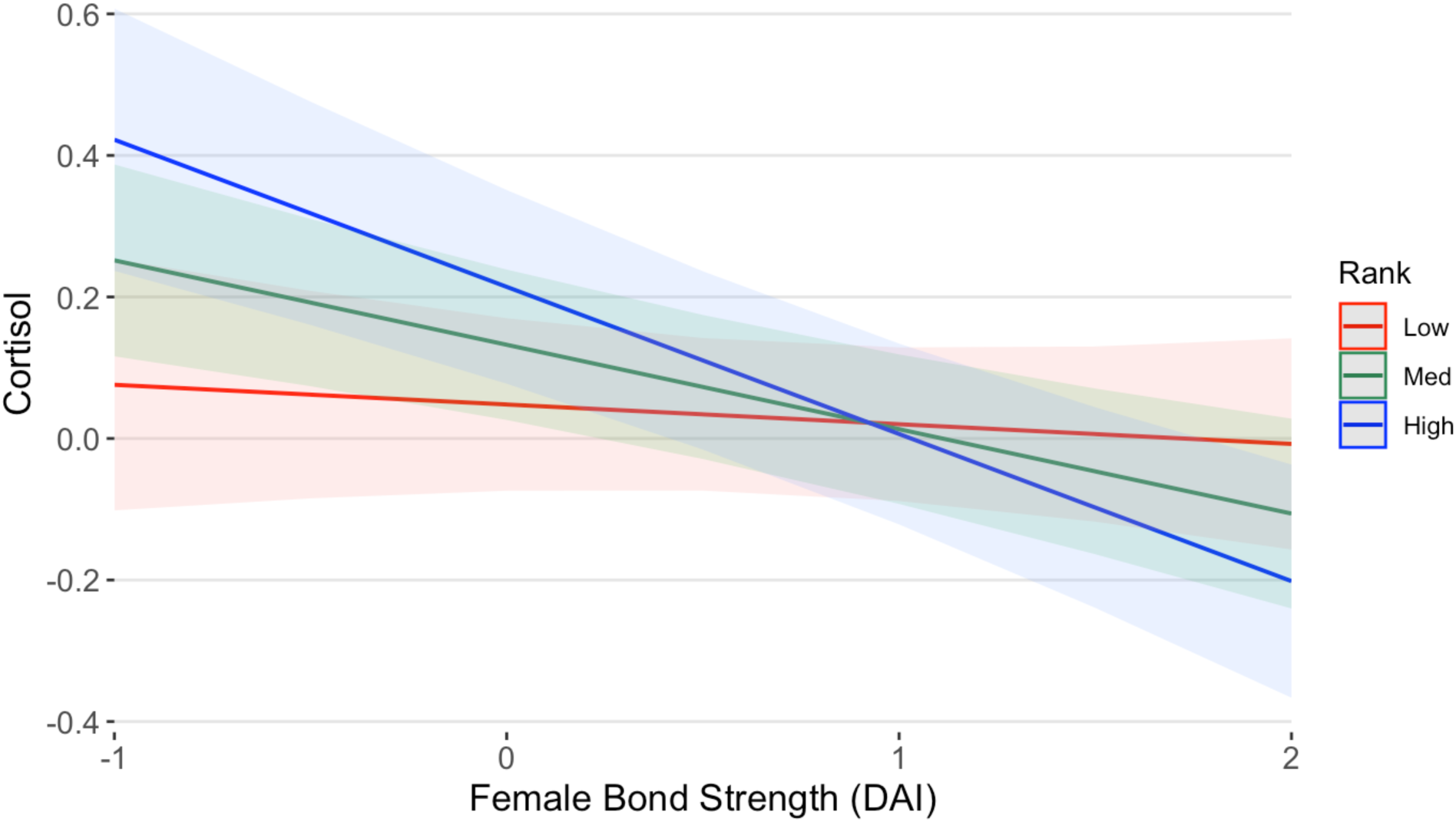
Effects of bond strength on female cortisol. Results of linear mixed-effects models showing marginal effects (±95% confidence intervals) of dominance rank and bond strength (Dyadic Association Index) on urinary cortisol levels in female chimpanzees. Cortisol values are standardized to time of day and specific gravity and log-transformed.

The effects of bonds were context-dependent in males, but none of the interactions we tested supported the hypothesis that bond strength reduces the impact of stressors. We found a significant positive interaction between instability and DSI (Figure 2a, est = 0.25, P = 0.022). This effect was in the opposite direction predicted, indicating that hierarchy instability raised cortisol levels more among strongly-bonded males than weakly-bonded males. We also detected a significant negative interaction between mating contexts and DSI in males (est = -0.28, P = 0.004). This interaction was in the direction predicted, such that availability of parous, swollen females increased the cortisol levels of weakly-bonded males more than strongly-bonded males, but the cortisol levels of strongly-bonded males remained consistently high regardless of mating context (Figure 2b). The interaction between DSI and dominance rank did not significantly improve model fit, so bonds did not buffer the costs of high rank.

**Figure 2.**
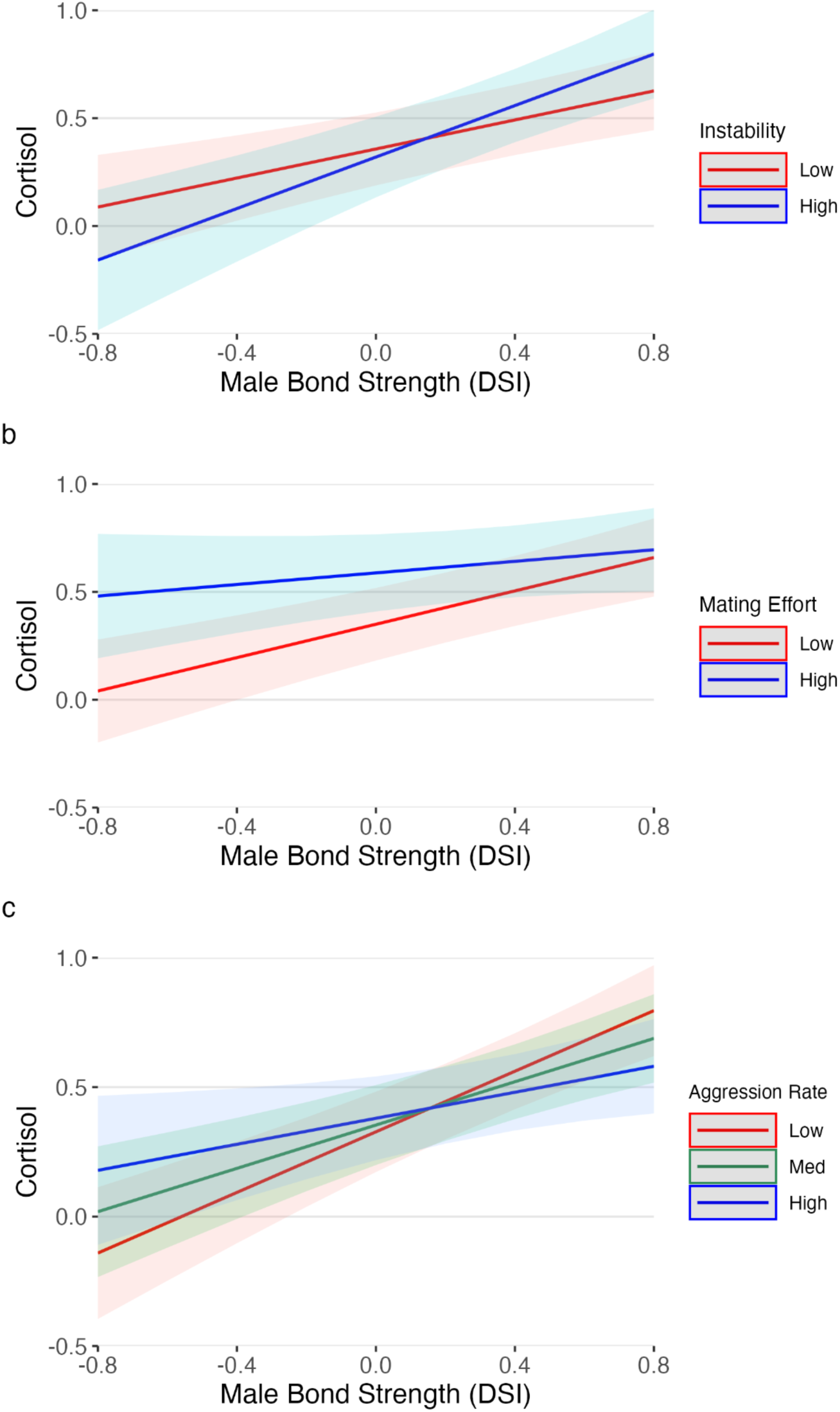
Effects of bond strength on male cortisol. Results of linear mixed-effects models showing marginal effects (±95% confidence intervals) of bond strength on male cortisol, split by (a) hierarchy stability, (b) mating effort, and (c) aggression given. Cortisol values are standardized to time of day and specific gravity and log-transformed.

### The effects of bond strength were not moderated by age (P3)

While age had a significant positive effect on cortisol for both males and females, model fits were not improved for either sex by including the interaction between age and bond strength.

### The effect of bond strength on cortisol was independent of aggression (P4)

Independent of the other variables in our model, aggression did not explain significant additional variation in female cortisol levels (Table 1) and thus did not mediate the effect of bond strength. Effect sizes for bond strength were identical in a model that included aggression received and a version of our original model that used a comparable sample (Tables 3, S1). The negative relationship between cortisol and bond strength, therefore, cannot be explained by reduced exposure to aggression.

While rates of both giving and receiving aggression were associated with increased male cortisol levels, the main effect of bond strength could not be explained by participation in aggression, as it remained strongly positive (see Tables 2, 3, S2). However, the effect of giving aggression varied with bond strength (Figure 2c, Est = -0.103, P = <0.001). Specifically, bond strength was positively correlated with cortisol at all levels of aggression given, but being highly aggressive was associated with a larger increase in cortisol for weakly-bonded males versus strongly-bonded males. This interaction replaced the context-specific interactions of bond strength with instability and mating context. That is, where hierarchy instability and mating contexts alter the relationship between bond strength and cortisol, this can be explained by changes in rates of male aggression in these contexts.

**Table 3:**
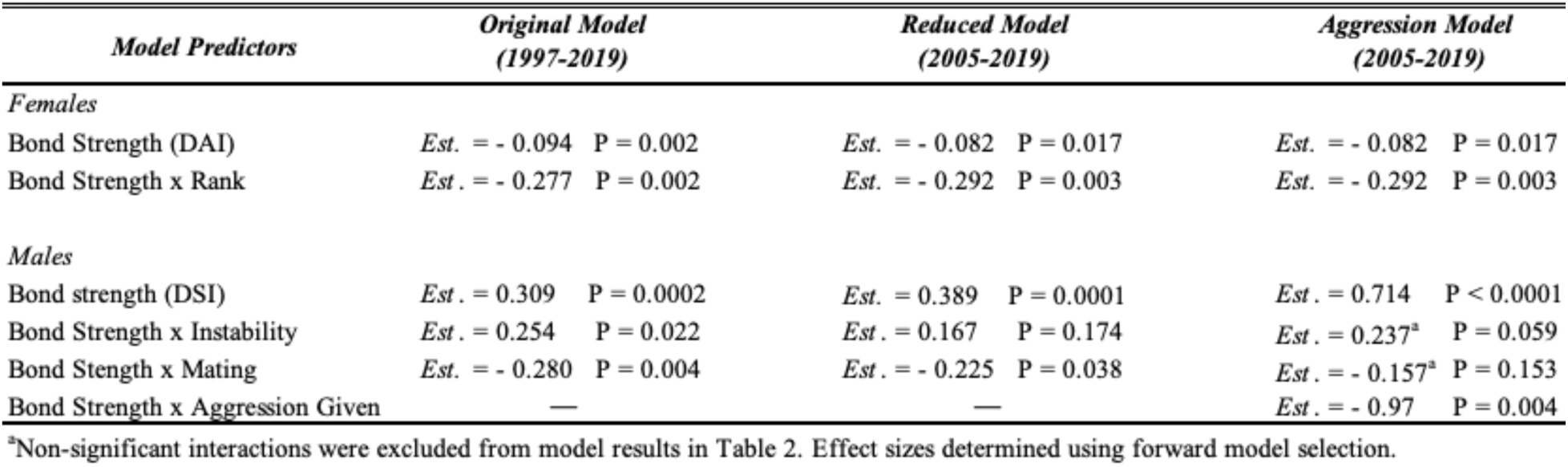
Comparison of effect sizes for bond strength across models with and without aggression. Results are from linear mixed-effects models testing the effects of bond strength on glucocorticoids. Primary models used the full dataset, while aggression models were fit to a reduced dataset with comprehensive aggression records. All estimates shown are for parameters significant in the aggression models.

## Discussion

We used a dense longitudinal dataset to test the hypothesis that strong social bonds reduce glucocorticoid production in wild chimpanzees. This hypothesis was supported for females, despite the fact that social bonds among female chimpanzees are generally weaker than among males [58,72]. In contrast, strongly-bonded males experienced higher cortisol levels than weakly-bonded males, and this relationship persisted across all social contexts we examined. While we found some context-dependent effects of social bonds, none of these results suggest that bonds were more protective under conditions when individuals experienced a higher rate of stressors. These effects also did not vary across age, despite increasing burden of high glucocorticoids in older chimpanzees [63]. While our previous findings indicated that rates of aggression fully mediated a strong association between high male rank and high glucocorticoids [36], aggression did not help explain the effects of social bonds on cortisol for either sex. These results introduce nuance to the wealth of evidence linking social support to positive health or fitness benefits in humans and other primates [2,8], highlighting a consideration that is rarely addressed: the formation and maintenance of social bonds can entail significant costs as well as benefits. Males improve their reproductive success through aggressive competition for rank [49,73], at least in part due to the metabolic cost of maintaining dominance [36]. Among male chimpanzees, social bonds are formed strategically to support these competitive efforts for status and mating opportunities [73], such that the long-term physiological costs of these relationships may be offset by their overall fitness advantages.

More broadly, the idea that sociality entails both benefits and costs has important biological implications: if participating in demanding social relationships increases physiological strain, individuals may differ in their capacity to engage in and sustain them. Such tradeoffs—between the fitness advantages of cooperation and its energetic costs—may help explain not only variation in male bonding strategies in chimpanzees, but also broader patterns of individual differences in social behaviour across species, including humans.

Social bonds among male chimpanzees yield more detectable fitness benefits. Males often leverage strong affiliative bonds for support in agonistic interactions, as well as to obtain mating concessions [45,47–50]. Thus, strong bonds, particularly those with well-placed individuals, are associated with higher rank acquisition and paternity success [49,74]. This dynamic of social bonding contrasts in important ways to what is typically observed in female mammals, where reproductive success must be accrued through longer periods of investment [75].

Other evidence on glucocorticoids and male social bonds is limited, likely because reproductive competition constrains affiliative relationships: in many species, males are intolerant of other mature males in their group [76]. Even in multimale primate groups, intense within-group competition for mates discourages strong social ties since, unlike food patches, fertilizations cannot be readily shared [77]. Nevertheless, in some species, males do form bonds, typically under moderate to low within-group contest competition [77]. These long-term alliances provide direct fitness benefits by supporting political coalitions, such as coalitionary support in dominance contests or collective defense. In two macaque species where the effects of bonds on glucocorticoids has been examined, social bonds helped buffer glucocorticoids responses to stressful contexts [22,27]. In human men, social support has been associated with buffered responses to stressful stimuli and overall more positive health outcomes, though notably the primary relationship in most of these studies is the marital bond or other long-term opposite-sex partnerships [78–80]. Male baboons bonded to females also experienced survival benefits [42]. Here, we did not examine intersexual bonds because most of the variation in affiliation between male and female chimpanzees is driven by the cycling status of females [59].

Previous reports on the health benefits of social bonds in non-human animals focus primarily on female-bonded species with strong matrilineal kin ties. Related females in these species cooperate to compete with other matrilines for food resources [55,81,82]. The inclusive fitness benefits of cooperating with kin are expected to minimize conflicts of interest over participation and division of payoffs [83]. Due to female-biased dispersal, most adult female chimpanzees lack access to close adult female kin [52]. Additionally, contest competition for food resources rarely occurs because high fission-fusion dynamics allow female chimpanzees to avoid competitors by exploiting the spatial distribution of food patches [84]. However, females in Kanyawara selectively tolerate a small number of other females, both close kin and non-kin, and they rely on these bonds on the rare occasions that they form coalitions [58,60]. Because access to high-quality core foraging areas confers substantial reproductive benefits, high-ranking females often defend these areas from newly immigrated females—one of the primary contexts in which female coalitions occur [85,86]. In this study, we found that social bonds more strongly reduced cortisol among high-ranking females. Thus, it is plausible that having a reliable bond partner reduces the costs of defending these ranges. Having a trusted relationship with a fellow mother may also facilitate infant socialization, as in orangutans [87,88], reducing the costs for mothers who are otherwise the primary play partners for their offspring [89]. Although mother–daughter dyads were rare and no sister dyads were present in our dataset, a robustness check excluding the few individual-years with kin was conducted to test whether kin relationships influenced our results. This did not change the main effect of strong social bonds on lower cortisol in females, indicating that this effect is not purely driven by kin relationships. These results are consistent with previous observations at Kanyawara, where female social bonds and cooperation are not solely determined by kinship [58,60]. Notably, after removing these kin-years, the effect of instability on cortisol became statistically significant and slightly stronger in non-kin contexts, consistent with the idea that close female kin can attenuate physiological responses to social instability. These findings suggest that the benefits of female social bonds are context-dependent and shaped by the limited availability of close partners. Our results raise the question of why, if advantageous, female social bonds are underutilized. In more recent years, several sisters and female offspring have remained in the natal community, and testing these cases could clarify how more abundant female kin networks influence cortisol responses to social instability.

A relatively straightforward explanation for the positive association between cortisol and bondedness might be that bonds are not costly, per se, but that males are more motivated to invest in social bonds when engaging in high-cost activities. Our findings do not support this view: males with strong bonds exhibited higher cortisol across all contexts, including those unrelated to dominance interactions, and this pattern persisted after controlling for aggression. This suggests that elevated cortisol is better explained by the demands of sustaining strategically important alliances than by dominance striving alone. In other words, for male chimpanzees, highly affiliative relationships appear, on average, to be costly in and of themselves. Several factors may contribute to this. First, males’ strongest allies in a group are also primary rivals for reproductive opportunities, meaning that maintaining bonds entails managing significant conflicts of interests [90]. High rates of affiliation among some dyads may follow from an increased need to reconcile conflicts. Similarly, some apparently strong relationships may result from a ‘keep your friends close and your enemies closer’ strategy, whereby males can monitor their rivals through close associations while using grooming to alleviate tensions between them. Second, the nature of male competition is such that the most effective allies are those with the most power or control over resources, rather than those with shared genetic interests [91]. Male chimpanzees often form bonds with non-kin [45,54], selecting partners based on traits such as rank, age, and competence, which increase the likelihood of effective coalitionary support [45], though evidence suggests they may also show a preference for maternal brothers [45,92]. However, although male association patterns at Kanyawara have been observed to be stable over many years [72], the composition and functional value of individual alliances can change as partners’ rank, age, or social leverage shifts, affecting access to the most valuable allies. Investments in high-value partners may go unreciprocated or even be met with aggression as individuals strategically navigate complex social hierarchies [93,94]. Navigating new bonds has indeed been associated with anxiety or glucocorticoid increases in humans [95] and juvenile primates [96,97]. Finally, cortisol responses to acute social stressors in humans are sometimes followed by increased affiliative behaviour [98], suggesting that glucocorticoids may motivate efforts to initiate or reinforce social connections. Thus, our results could be explained if males who perceive greater risk in their environment invest disproportionately in developing bonds, perhaps at the expense of more diverse social networks.

Social bonds could influence glucocorticoids either by impacting the rate at which individuals experience stressors or the ways in which individuals perceive or respond to those stressors. As a preliminary way to address this distinction, we assessed whether any effects of social bonds were mediated by aggression, the most obvious and frequent proximate stressor in chimpanzees. Overall, this was not the case, as including aggression in our model mediated neither the positive effect of social bonds on cortisol in males nor the negative effect in females. However, our results also do not provide clear support for a stress response buffering effect because strongly-bonded individuals were not more resistant to the cortisol increases that accompanied stressful contexts, namely hierarchy instability and mating competition. We note that our study does not present a clear test between these mechanisms because we were not measuring timed responses to controlled stressors. For example, while instability and mating effort are associated with group-wide increases in aggression and cortisol, the direct involvement of different individuals in those contexts likely differs. Indeed, where we observed context-specific effects of social bonds on cortisol in males, those interactions did not persist in a model that controlled for aggression. While there is evidence that social affiliation reduces the immediate cortisol response to a stressor in wild chimpanzees [23], these naturalistic studies similarly lack the ability to control the intensity of stressful stimuli experienced, or confounding effects of unmeasured stressors.

There are prior reports that at least some forms of social bonds carry significant costs. In facultatively social yellow-bellied marmots (*Marmota flaviventris)*, females with stronger affiliative relationships died at younger ages [99]. In female blue monkeys, strong female bonds were associated with higher survival if those bonds were stable, but reduced survival when there was a high turnover between partners [100]. Among mountain gorillas (*Gorilla beringei beringei*), strong and stable social bonds were associated with reduced injury risk but increased illness in males, and with positive or negative effects in females that depended on group size [101]. In humans, the marital bond is typically associated with improved health and survival, but people in distressed marriages experience worse outcomes than those who were never married [102,103]. Individuals who take on a significant caregiving role in a close relationship with a friend or relative also experience a high burden of stress and accelerated aging [104,105]. Taken together, this literature suggests that investment in social bonds can be associated with risks and conflicts of interests that, if not successfully managed, produce negative short- or long-term consequences. Where individuals must pay significant costs to reap fitness rewards from social bonds, this will also be an important constraint on social participation. Such costs could help explain the highly selective engagement in social bonds from chimpanzee females, both in this community and across other populations [60]. And while we did not find evidence that social bonds differentially influenced cortisol among older chimpanzees, older or less healthy individuals may still be expected to reduce social investments if they are less able to tolerate these costs. Prior results from chimpanzees, humans, and other primates indicate significant reductions in social effort with age, though often with an increase in an investment in the most reliable bond partners (chimpanzees: [106,107]; humans:[108]; other primates: [109–111]).

Chronic activation of the HPA axis has well-documented links to poor health and reduced survival across species [2,14], including in wild baboons, where higher glucocorticoid levels predicted reduced adult female and offspring survival [41]. Social stress in response to status competition has also been linked to accelerated aging in baboons [13] and rhesus macaques [112]. We identified strong links between social environments and glucocorticoids that appear to persist beyond the immediate response to stressors. We are cautious in our interpretation, since we have not yet determined whether these links are associated with health outcomes [113]. Conservatively, we can conclude that high glucocorticoid levels associated with some chimpanzee social environments are costly, evidence of metabolic responses to real or perceived demands. We may expect that repeated exposure to these demands imposes constraints on key life-history processes, including growth, reproduction, and aging.

## Ethics

This research received approval from the Uganda National Council for Science and Technology, the Uganda Wildlife Authority, and the Makerere University Biological Field Station. Our study protocols were reviewed and approved by the Institutional Animal Care and Use Committees at the University of New Mexico, Tufts University, and Harvard University.

## Supporting information

READ ME file for supplement

male dataset

female dataset

female dataset excluding kin bonds

Analysis R code

## Data accessibility

The data supporting this article are provided as part of the electronic supplementary material.

