## Supplementary material for "Sex-specific effects of social bonds on glucocorticoids in wild chimpanzees": READ ME file for supplement

README: Analysis of Social Bonds on Cortisol in Wild Chimpanzees

========================================================

Title of Study:

---------------

Sex-specific effects of social bonds on glucocorticoids in wild chimpanzees

Short Summary:

--------------

This repository contains the R code and data used to analyze the relationship

between social bond strength and stress hormone (cortisol) levels in wild

chimpanzees. Analyses include linear mixed-effects models and figures summarizing

predicted cortisol levels.

Repository Contents:

-------------------

- analysis_code.md	: Main R code for all analyses, including data processing,

model fitting, and figure generation.

- Female_data.csv	: Anonymized dataset for female chimpanzees. Contains

all variables used in models and figures.

- Male_data.csv	: Anonymized dataset for male chimpanzees. Contains

all variables used in models and figures.

- Female_data_nokin.csv : Subset of female data excluding mother-daughter kin dyads

for kinship bias checks.

- figpub_females.png : Figure summarizing predicted cortisol for females (Figure 1).

- figpubs_males.png : Figures summarizing predicted cortisol for males (Figure 2, combined).

- README.txt	: This file.

\## Data: Column Headers and Units

\### Female_data.csv

| Column | Description | Units / Notes |

| ChimpID | Unique identifier for each female | - |

| Year | Calendar year of sample | yyyy |

| YearMonth | Year and month of sample | yyyy-mm |

| Age | Age of female at sampling | years; centered on 15 |

| Rank | Dominance rank | ordinal (low to high); centered on 0.5; same-sex hierarchy |

| ReproStatus | Reproductive status | categorical; 4 levels: cycling w/ max swelling, cycling w/o max swelling, early lactation, late lactation; reflects mating effort |

| Instability | Social instability index | 0/1 per date; 1 if any reversals in relative ranks of two or more females in previous 4 weeks |

| DAI | Dyadic association index (bond strength) | unitless; centered on 1 |

| GroomingTop3 | Grooming interactions with top 3 partners| count |

| Cortisol | Cortisol concentration | standardized log-transformed |

| VictimRate | Rate of received aggression | events per hour per individual; corrected for observation time |

\### Female_data_nokin.csv

| Column | Description | Units / Notes |

| ChimpID | Unique identifier for each female | - |

| Year | Calendar year of sample | yyyy |

| YearMonth | Year and month of sample | yyyy-mm |

| Age | Age of female at sampling | years; centered on 15 |

| Rank | Dominance rank | ordinal (low to high); centered on 0.5; same-sex hierarchy |

| ReproStatus | Reproductive status | categorical; 4 levels: cycling w/ max swelling, cycling w/o max swelling, early lactation, late lactation; reflects mating effort |

| Instability | Social instability index | 0/1 per date; 1 if any reversals in relative ranks of two or more females in previous 4 weeks |

| DAI | Dyadic association index (bond strength) | unitless; centered on 1 |

| GroomingTop3 | Grooming interactions with top 3 partners| count |

| Cortisol | Cortisol concentration | standardized log-transformed |

| VictimRate | Rate of received aggression | events per hour per individual; corrected for observation time |

\### Male_data.csv

| Column | Description | Units / Notes |

| ChimpID | Unique identifier for each male | - |

| Year | Calendar year of sample | yyyy |

| YearMonth | Year and month of sample | yyyy-mm |

| Age | Age of male at sampling | years; centered on 15 |

| Rank | Dominance rank | ordinal (low to high); centered on 0.5; same-sex hierarchy |

| MatingEffort | Effort toward mating | binary (0/1); at least one parous, maximally-swollen female observed in party; reflects mating effort|

| Instability | Social instability index | 0/1 per date; 1 if any reversals in relative ranks of two or more males in previous 4 weeks |

| DSI | Dyadic sociality index (bond strength) | unitless; centered on 1 |

| AggressionRate | Rate of aggression initiated | log-transformed; events per hour per individual; corrected for observation time |

| Cortisol | Cortisol concentration | standardized log-transformed |

| VictimRate | Rate of received aggression | events per hour per individual; corrected for observation time |

**Notes on transformations and centering:**

- Where appropriate, fixed effects were log transformed (AggressionRate) or centered on a meaningful zero value (Age = 15, Rank = 0.5, Bond strength = 1).

- Aggression: recorded using all-occurrence sampling; events between same victim-aggressor counted only if ≥10 min apart.

- Dominance ranks: determined by daily Elo ratings. All ranks were standardized for the number of individuals in the hierarchy, so that the lowest-ranking individual of each sex received a score of 0, while the highest-ranking individual received a score of 1.

- Instability: 0/1 per date, reflecting rank reversals in previous 4 weeks.

- Female ReproStatus: covariate for reproductive status (4 levels) reflecting mating effort.

- Male MatingEffort: binary (0/1), reflects presence of at least one parous, maximally-swollen female.

- AggressionRate and VictimRate: per-hour rate per individual, corrected for observation hours.

Code and Package Versions:

--------------------------

- R version: 4.5.1

- Packages used: dplyr, tidyverse, readr, lme4, lmerTest, sjPlot, ggplot2,

grid, gridExtra, cowplot

- Session info included at the end of analysis_code.md for reproducibility.

How to Run the Code:

--------------------

1. Open analysis_code.md in RStudio.

2. Ensure all CSV files are in the same working directory as the Markdown file.

3. Install required packages if not already installed:

install.packages(c("dplyr", "tidyverse", "readr", "lme4", "lmerTest",

"sjPlot", "ggplot2", "grid", "gridExtra", "cowplot"))

4. Run the code chunks sequentially. Figures will be generated in the working directory.

Mapping of Code to Output:

--------------------------

- Sections 3–5: Fit linear mixed-effects models for females and males

(including aggression and kinship subset checks). Models correspond to Tables 1–3 and S1–S2.

- Section 7: Generate figures from model predictions.

- figpub_females.png: Female cortisol predictions

- figpubs_males.png: Male cortisol predictions (combined a–c panels)

- Section 8: Session info for reproducibility

Notes on Data:

--------------

- All data are anonymized; no chimpanzee identifiers are included.

- CSV files represent the most complete version of each dataset (prior to any filtering).

- Figures are provided as .png images to match the manuscript. Source data for figures

are included in the CSVs.

Contact:

--------

- Initial submission is blinded, so author info is omitted here.

- For questions post-publication, contact: []
