## Supplemental Text and Tables for "Sex-specific effects of social bonds on glucocorticoids in wild chimpanzees"

### Methods

We analyzed variation in urinary glucocorticoid concentrations in adult chimpanzees (aged 15 years and older) in the Kanyawara community of wild, unprovisioned chimpanzees in Kibale National Park, Uganda, between Dec 1997 and Dec 2019. During this study period, the community size ranged from 41-56 individuals. There were 9-13 adult males and 13-18 adult females in any given study year. Prior to any exclusions, as noted below, we assayed 11034 samples from females and 9875 for males.

*Cortisol*

Urine was captured either by placing plastic in the stream as the chimpanzees urinated from the trees or by pipetting from forest vegetation. We collected only when the identity of the urinating chimpanzee could be confirmed and the site was uncontaminated by feces or by the urine of other individuals. Samples were stored on ice until the end of the observation day, and any particulates were removed prior to freezing at -20 degrees C.

Immunoreactive cortisol was determined in urine samples via enzyme immunoassay with reagents and protocols provided by the University of California, Davis Clinical Endocrinology Laboratory (antibody R4866). Specifics of the protocols and longitudinal performance validation characteristics are described elsewhere [1]. Prior to analysis, raw cortisol concentrations were transformed in three ways. First, concentrations were standardized to the population mean specific gravity of 1.017 to correct for variation in urine water content. Second, the resulting standardized concentrations were log-transformed to approach normality. Third, we adjusted for diurnal variation in cortisol excretion by deriving residuals from a cubic spline regression for time of excretion [2,3]. The unit of analysis was the time-adjusted residual cortisol of each urine sample. Although the assay demonstrates strong interassay reliability [3], we additionally included in the time correction model categorical control variables for (a) the year in which the assay was performed in the laboratory, and (b) storage time before assays, defined categorically (< 4 years, 61% of samples; 4-6 years, 21% of samples; 7-9 years, 18% of samples). The latter addresses a trend discovered in exploratory analyses, where storage time correlates with a minor, but statistically significant and non-linear decline in measured cortisol concentrations . Samples were excluded if they were overly dilute, with specific gravity < 1.003, or if they were outliers with cortisol concentrations > 2 SD from the mean. We also excluded pregnant females from the study, as glucocorticoids are elevated during pregnancy due to altered adrenal function [4]. Four females with one or more urine samples were excluded due to insufficient behavioral data.

**Behavioral data collection**

Social behavior was observed by teams of trained observers, typically comprising at least 2-3 individuals collaborating to observe a social group from multiple angles [5]. Throughout the study, party membership was recorded at 15-minute intervals using scan sampling. A chimpanzee party includes all individuals who are spaced no more than 50m from another party member. We attempted to follow chimpanzees every day for the entire active period, and multiple observation teams often collected data on separate parties. A total of 89,913 hours of group observation contributed to this study.

Proximity and grooming data were derived from focal observations. Between December 1997 and July 2009, focals were ten minutes in duration with proximity and grooming recorded every two minutes. To preserve independence, we included only the final scan, i.e., once per 10 minutes. From Aug 2009 to Dec 2019, we conducted full-day focals of the same individual whenever possible, switching to another individual only if the original focal was unavoidably lost. Proximity was recorded every 15 minutes throughout the focal, and all occurrences of grooming behaviour involving the focal were recorded. Behavioural measures from each dataset were converted to rates based on the appropriate denominator for sampling interval, and as below, data were standardized to year-specific means. An average of 829 hours of focal observation were collected per individual (range: 5-2823 hrs) for 23 males and 28 females.

Grooming was assessed dyadically by the frequency of bouts between two partners. A bout could include sequential or simultaneous grooming by each partner and was defined as an independent bout if separated by at least 10 minutes from another bout between the same dyad.

Aggression was recorded using all-occurrences sampling [2] and defined as a rate per individual, corrected for the number of hours observed. Events between the same victim and aggressor were only counted as independent events if separated by at least 10 minutes. Dominance ranks were determined in same-sex hierarchies based on decided, dyadic agonistic interactions and pant-grunts, formal signals of submission [2,3].

All male ranks are based on daily Elo ratings [6]. Female ranks are much more stable, but interactions less frequent, than in males [7,8]. Accordingly, the Elo method was unable to resolve female hierarchies before 2004 due to sparseness of interactions, so earlier female rankings were determined by calculating Modified David's scores on interaction matrices [3,9].

**Bond strength**

To characterize the bondedness of adult males, we calculated the dyadic composite sociality index (DSI), following methods similar to those used elsewhere in primates [10]. The measure averages three component behavioral indices (party association, proximity, grooming) that are each standardized for the mean rate of interactions across all dyads of the same sex composition in the same year (average DSI = 1). Because chimpanzees have high fission-fusion dynamics, our DSI includes a party association index not typically necessary for other primates. Party association rate was calculated from all group observations as the number of 15-minute group scans in which both individuals were present in the same party divided by the total number of scans in which at least one of those individuals was observed. Proximity rate was calculated from focal observations as the number of scans (10 or 15 minute intervals) in which the dyads members were within 5 meters of one another as a proportion of the total number of scans in which (a) the dyad was present together in the same party; and (b) one of them was the focal. The proximity index was only included where the dyad members were observed together for at least 6 hours or followed as individual focals for a combined 20 hrs. The last criteria allows us to reduce sampling bias (partners were not observed interacting because there were too few opportunities to do so) while including true zeroes (partners had sufficient opportunity to interact but did not). Grooming rate was calculated from focal observations as the number of grooming bouts between a dyad relative to the number of scans where they were within 5m of one another while one was the focal. The grooming index was only calculated when the dyad spent at least 1 hour in proximity. While preferred partners are likely to have high rates of interaction across all measures, each successive measure in the index applies a different denominator for statistical independence from the prior measure. To ensure that all dyads had ample opportunity to interact, we only included those where (a) each chimpanzee was observed for at least 100 hours during the year, and (b) dyad members were simultaneously present in the community as adults for at least 6 months of that year (i.e., had ample opportunity to interact). 235 (10.9%) of the annual male dyads did not meet the minimum criteria to calculate DSI. These can be assumed to be those who were not bonded, and therefore these exclusions should not affect our results. *Bond strength* for males was defined as the average of the highest 3 DSIs (strongest 3 dyads) for each male in each year.

In this community, grooming between female chimpanzees is rare, and recent studies indicate that shared proximity better explains tolerance and cooperation among partners than rate of grooming [11,12]. The DSI measure was not suitable for females because a large proportion of dyads failed to meet the minimum rate of proximity necessary to compute the grooming index, and grooming was rare. Indeed, 84% of the annual female-female dyads in our study did not groom at all, and only 3% of female-female dyads groomed more than 3 times in a year. Accordingly, for the female analyses, we substituted a dyadic association index (DAI), which was identical to the measure for males but without the grooming component. Thus, *bond strength* for females was defined as the average of the highest 3 DAIs for each year. Separately, we included a covariate for *grooming* which indicated how many of these top 3 social partners the female had groomed with at all during the year (range 0-3). Note that in some years, females did not meet the minimum inclusion criteria to calculate DAI for at least 3 partners, and datapoints for these females were excluded for the full year, reducing our sample by 17%.

We did not consider opposite sex affiliative relationships in this analysis. Prior studies of this community indicate that, while female chimpanzees associate and groom with males at moderately high rates, these relationships are temporary and based on the sexual receptivity of females [13].

**Data analysis**

To test the study predictions, we performed linear mixed models using each sample as the unit of analysis. Model structure was based on two prior analyses of a previous version of this longitudinal dataset [2,3] with newly-specific fixed effects to quantify the strength of same-sex affiliative bonds (for males = DSI, for females = DAI and Grooming), calculated for each individual in each calendar year.

Control variables were those which were previously found to have significant influences on cortisol in these prior studies and were specified in the same manner. We summarize these here, and refer readers to the previous studies for more detail.

· *Age* in years on day of sampling. Age was previously found to have a strong positive effect on cortisol in both sexes [2,3].

· Dominance *Rank* in the same-sex hierarchy on the day of sampling. All ranks were standardized for the number of individuals in the hierarchy, such that the lowest-ranking individual of each sex received a score of 0 while the highest-ranking individual received a score of 1. Rank was previously found to have a strong positive effect on cortisol in males but did not show a strong effect in females [2,3].

· Group *Instability*, defined as 0/1 on each date, based on whether there were any reversals in the relative ranks of two or more males in the previous 4 weeks. Instability at the group level was previously found to drive increases in male cortisol across the group, regardless of whether the individual male's own rank was affected [2].

· *Mating* Context. For females, we included a covariate for reproductive status using 4 levels: cycling females with maximally tumescent swellings, cycling females without maximally tumescent swellings, females in early lactation (first 2 years postpartum), or females in late lactation (2 years postpartum to resumption of cycling). Cycling females with maximal swellings have previously been found to have significantly higher cortisol than the three other categories. For males, we defined mating context using a binary variable (0/1) as whether at least one parous, maximally-swollen female was observed in a party with the sampled male on the day of sampling, as this has previously been associated with substantial elevations in male cortisol [2,3].

After performing our primary analysis to detect the effects of bond strength on cortisol, we conducted a secondary analysis to determine whether the effect of bond strength was mediated by variation in *aggression received*, and for males, *aggression given*. We added a fixed effect for rate of aggression received in a 14 day window prior to and including the date of sampling and additionally examined a potential interaction between aggression received and bond strength. Comprehensive aggression data were available from 2005 forward, and aggression rates were only considered if the chimpanzee was observed for at least 20 hours in that 14 day window, so these secondary analyses were based on a smaller subset of samples. To test for mediation, we compared effect sizes from this model with a similarly-reduced dataset that did not include aggression.

All models included random intercepts for chimpanzee ID and month-year of sample collection to minimize the influence of uneven sampling and unexplained temporal or variation. We considered random slopes for rank and age, but due to convergence or singularity issues, neither was retained in the female models and only the slope for rank was included in the male models. Because we had strong *a priori* reasons for including all fixed effects, all were retained in final models. We considered interactions of rank and age, bond strength and age, bond strength and rank, bond strength and instability, bond strength and mating context, and bond strength and aggression, and we only retained these when inclusion of the interaction significantly improved the model fit as assessed with a log-likelihood ratio test.

### Supplementary Results

We tested whether rates of aggression mediate the observed effects of social bonds on urinary cortisol. Tables 1, 2 and 3 in the main manuscript provide the results of the full cortisol datasets without aggression (Tables 1, 2) and comparison model including aggression (Table 3). The comparison model for aggression was based on a reduced sample including 6730 samples for 23 adult females (177 female-years) and 7086 samples from 17 adult males (140 male-years). In order to conduct a valid mediation, effect sizes from the aggression models were compared to a reduced version of the original model, as reported in Tables S1 and S2.

**Robustness to kinship:**

As an additional robustness check, we removed female-years in which individuals had a mother or daughter present in the dataset. Following this exclusion, the effect of hierarchy instability on cortisol became statistically significant (**estimate =** 0.071, $p = 0.025$), demonstrating that instability has a measurable positive effect on cortisol in non-kin contexts. Importantly, this exclusion did not alter the main effect of bond strength or the instability × bond strength interaction in either the primary or aggression comparison models, indicating that our primary results are robust to the presence of close kin.

This pattern likely reflects a buffering effect of mother–daughter relationships: in years when females had a mother or daughter present, the physiological effects of social instability were partially mitigated. Effect sizes in the kin-excluded dataset were slightly higher than in the full dataset, but the overall direction and magnitude of effects remained consistent.

**Table S1: Primary female model (aggression data subset).**

Full parameter estimates from the primary female linear mixed model using only samples with comprehensive aggression data. Results were compared to models including aggression and summarized in Table 3.

| | **Primary Model** | | | | | |
|---|---|---|---|---|---|---|
| *Model Predictors* | *Estimates* | *Variance* | *SE* | *SD* | *t* | *P* |
| Intercept | 0.011 | | 0.069 | | 0.160 | 0.872 |
| Age | **0.010** | | **0.003** | | **9.52** | **0.006** |
| Rank | **0.268** | | **0.128** | | **2.10** | **0.0397** |
| Instability | **0.068** | | **0.033** | | **2.06** | **0.039** |
| Mating: cycling + maximal swelling | **0.263** | | **0.054** | | **4.86** | **<0.001** |
| Mating: Early lactation | -0.033 | | 0.038 | | -0.87 | 0.385 |
| Mating: late lactation | -0.075 | | 0.044 | | -1.71 | 0.087 |
| Bond strength (DAI) | **-0.082** | | **0.034** | | **-2.39** | **0.017** |
| DAI x Rank | **-0.292** | | **0.097** | | **-3.02** | **0.003** |
| *Random Effects* | | | | | | |
| Chimp ID | | 0.020 | | 0.142 | | |
| Month/Year of sample collection | | 0.185 | | 0.430 | | |

N = 6730 urine samples, 177 female-years, 23 adult females
Random-effects variance components > 0 are interpreted as meaningful contributions of the grouping factor.

**Table S2: Primary male model (aggression data subset).**

Full parameter estimates from the primary male linear mixed model using only samples with comprehensive aggression data. Results were compared to models including aggression and summarized in Table 3.

| | **Primary Model** | | | | | |
|---|---|---|---|---|---|---|
| *Model Predictors* | *Estimates* | *Variance* | *SE* | *SD* | *t* | *P* |
| Intercept | **-0.153** | | **0.063** | | **-2.42** | **0.021** |
| Age | 0.002 | | 0.003 | | 0.88 | 0.389 |
| Rank | 0.234 | | 0.231 | | 1.01 | 0.329 |
| Instability | 0.047 | | 0.053 | | 0.88 | 0.380 |
| Mating: presence of swollen females | **0.241** | | **0.046** | | **5.27** | **<0.001** |
| Bond strength (DSI) | **0.387** | | **0.099** | | **3.90** | **<0.001** |
| DSI x Instability | 0.167 | | 0.123 | | 1.36 | 0.174 |
| DSI x Mating | **-0.225** | | **0.108** | | **-2.08** | **0.038** |
| Age x Rank | -0.005 | | 0.014 | | -0.39 | 0.698 |
| *Random Effects* | | | | | | |
| Chimp ID | | 0.022 | | 0.148 | | |
| Month/Year of sample collection | | 0.141 | | 0.376 | | |

N = 7086 urine samples, 211 male-years, 17 adult males
Random-effects variance components > 0 are interpreted as meaningful contributions of the grouping factor.

**Table S3: Female aggression model with full covariates (aggression data subset).**
Full parameter estimates from the secondary female linear mixed model including aggression variables (aggression given and received), restricted to samples with comprehensive aggression data. These results complement the reduced primary model (Table S1) and summaries for comparison are in Table 3.

| Aggression Model | | | | | | |
|---|---|---|---|---|---|---|
| *Model Predictors* | *Estimates* | *Variance* | *SE* | *SD* | *t* | *P* |
| Intercept | -0.030 | | 0.074 | | -0.411 | 0.682 |
| Age | **0.010** | | **0.003** | | **2.96** | **0.006** |
| Rank | **0.266** | | **0.128** | | **2.08** | **0.041** |
| Instability | **0.068** | | **0.033** | | **2.04** | **0.042** |
| Mating: cycling + maximal swelling | **0.258** | | **0.054** | | **4.77** | **<0.001** |
| Mating: Early lactation | -0.022 | | 0.039 | | -0.57 | 0.568 |
| Mating: late lactation | -0.066 | | 0.044 | | -1.49 | 0.135 |
| Bond strength (DAI) | **-0.292** | | **0.034** | | **-2.39** | **0.017** |
| DAI x Rank | **-0.292** | | **0.097** | | **-3.02** | **0.003** |
| Aggression Received | 0.012 | | 0.007 | | 1.61 | 0.107 |
| *Random Effects* | | | | | | |
| Chimp ID | | 0.020 | | 0.142 | | |
| Month/Year of sample collection | | 0.184 | | 0.429 | | |

N = 6730 urine samples, 177 female-years, 23 adult females
Random-effects variance components > 0 are interpreted as meaningful contributions of the grouping factor.

**Table S4: Male aggression model with full covariates (aggression data subset).**
Full parameter estimates from the secondary male linear mixed model including aggression variables (aggression given and received), restricted to samples with comprehensive aggression data. These results complement the reduced primary model (Table S2) and summaries for comparison are in Table 3.

| Aggression Model | | | | | | |
|---|---|---|---|---|---|---|
| *Model Predictors* | *Estimates* | *Variance* | *SE* | *SD* | *t* | *P* |
| Intercept | **-0.255** | | **0.073** | | **-3.50** | **<0.001** |
| Age | 0.002 | | 0.003 | | 0.71 | 0.485 |
| Rank | 0.273 | | 0.232 | | 1.17 | 0.259 |
| Instability | 0.015 | | 0.054 | | 0.28 | 0.779 |
| Mating: presence of swollen females | **0.219** | | **0.046** | | **4.77** | **<0.001** |
| Bond strength (DSI) | **0.726** | | **0.133** | | **5.45** | **<0.001** |
| DSI x Instability | 0.247 | | 0.125 | | 1.97 | **0.049** |
| DSI x Mating | -0.168 | | 0.109 | | -1.53 | 0.126 |
| Age x Rank | -0.004 | | 0.014 | | -0.31 | 0.760 |
| Aggression Received | **0.019** | | **0.007** | | **2.71** | **0.007** |
| Aggression Given | 0.019 | | 0.011 | | 1.65 | 0.099 |
| DSI x Aggression Given | **-0.102** | | **0.028** | | **-3.59** | **<0.001** |
| *Random Effects* | | | | | | |
| Chimp ID | | 0.021 | | 0.144 | | |
| Month/Year of sample collection | | 0.140 | | 0.374 | | |

N = 7086 urine samples, 211 male-years, 17 adult males
Random-effects variance components > 0 are interpreted as meaningful contributions of the grouping factor.

### Data Accessibility

See separate files for full datasets, R code, and README file for instructions on statistical replication.
