## Supplementary material for "Sex-specific effects of social bonds on glucocorticoids in wild chimpanzees": Analysis R code

\# -------------------------------

\# Analysis Code

\# -------------------------------

This file contains the R code used to generate analyses and figures for the study.

---

```{R, echo=FALSE}

\# -------------------------------

\# 1. Setup

\# -------------------------------

\# Load required packages

library(dplyr)

library(tidyverse)

library(readr)

library(lme4)

library(lmerTest)

library(sjPlot)

library(ggplot2)

library(grid)

library(gridExtra)

library(cowplot)

\# -------------------------------

\# Global R Markdown chunk options

\# -------------------------------

knitr::opts_chunk$set(

comment = NA, # remove default "#>" before output

message = FALSE, # hide package loading messages

warning = FALSE, # hide warnings

echo = TRUE, # show code in the .md submission

fig.align = "center",# center all figures

fig.width = 6, # default figure width (inches)

fig.height = 4, # default figure height (inches)

dpi = 300, # high resolution for publication figures

fig.path = "figures/", # optional: save figures in a subfolder

out.width = "80%", # scale figure width in output

out.height = "auto", # keep aspect ratio

tidy = TRUE	# cleanly format code in output

)

\# -------------------------------

\# 2. Data Import (Datasets provided in supplement)

\# -------------------------------

\### All datasets used in these analyses are provided as CSV files in the Supplementary Materials

\## 2a. Female data

\# CSV file: Female_data.csv (Supplementary Material)

Female_data <- read.csv("Female_data.csv")

\## 2b. Male data

\# CSV file: Male_data.csv (Supplementary Material)

Male_data <- read.csv("Male_data.csv")

\# -------------------------------

\# 3. Primary Models (Full Dataset)

\# -------------------------------

```
### 3a. Females preliminary model (excluding aggression)
### Results shown in Tables 1, 3
```

Females <- lmer(

Cortisol ~ Age + Rank + ReproStatus + Instability + DAI + Rank*DAI +

(1|ChimpID) + (1|YearMonth),

na.action = na.omit,

data = Female_data

)

```
### 3b. Males preliminary model (excluding aggression)
### Results shown in Tables 2, 3
```

Males <- lmer(

Cortisol ~ Age + Rank + Age*Rank + MatingEffort + Instability + DSI + DSI*Instability + DSI*MatingEffort +

(1+Rank|ChimpID) + (1|YearMonth),

na.action = na.omit,

data = Male_data

)

\# -------------------------------

\# 4. Aggression: Data Subset and Models

\# -------------------------------

```
### 4a. Females: only rows with VictimRate
### Results shown in Tables 3
```

Female_data_agg <- Female_data %>% filter(!is.na(VictimRate))

Females_agg <- lmer(

Cortisol ~ Age + Rank + ReproStatus + Instability + DAI + Rank*DAI + VictimRate +

(1|ChimpID) + (1|YearMonth),

na.action = na.omit,

data = Female_data_agg

)

```
### 4b. Males: only rows with VictimRate & AggressionRate
### Results shown in Tables 3
```

Male_data_agg <- Male_data %>% filter(!is.na(VictimRate) & !is.na(AggressionRate))

Males_Agg <- lmer(

Cortisol ~ Age + Rank + Age*Rank + MatingEffort + Instability + DSI + DSI*Instability + DSI*MatingEffort +

VictimRate + AggressionRate + AggressionRate*DSI +

(1+Rank|ChimpID) + (1|YearMonth),

na.action = na.omit,

data = Male_data_agg

)

\# -------------------------------

\# 5. Primary Models on Aggression Subset (for comparison)

\# -------------------------------

```
### 5a. Females
### Results shown in Table S1
```

Females_agg_comparison <- lmer(

Cortisol ~ Age + Rank + ReproStatus + Instability + DAI + Rank*DAI +

(1|ChimpID) + (1|YearMonth),

na.action = na.omit,

data = Female_data_agg

)

```
### 5b. Males
### Results shown in Table S2
```

Males_agg_comparison <- lmer(

Cortisol ~ Age + Rank + Age*Rank + MatingEffort + Instability + DSI + DSI*Instability + DSI*MatingEffort +

(1+Rank|ChimpID) + (1|YearMonth),

na.action = na.omit,

data = Male_data_agg

)

\# -------------------------------

\# 6. Kinship Bias Check (Females Only)

\# -------------------------------

```
### No male kin dyads observed in dataset
### Dataset excludes samples in years when female had a kin dyad
```

\# CSV file: Female_data_nokin.csv (Supplementary Material)

#Female_data_nokin <- read.csv("data/Female_data_nokin.csv") # path relative to project folder

Female_data_nokin <- read.csv("/Users/ellendyer/Desktop/PROCEEDINGSB/Female_data_nokin.csv")

\## Check number of samples removed

n_removed <- nrow(Female_data) - nrow(Female_data_nokin)

n_removed

\## Fit model without kin samples

Fem_model_nokin <- lmer(

Cortisol ~ Age + Rank + ReproStatus + Instability + DAI + Rank*DAI +

(1|ChimpID) + (1|YearMonth),

na.action = na.omit,

data = Female_data_nokin

)

\## Check whether interaction between bond strength x instability becomes significant

Fem_model_nokin1 <- lmer(

Cortisol ~ Age + Rank + ReproStatus + Instability + DAI + Rank*DAI + Instability*DAI +

(1|ChimpID) + (1|YearMonth),

data = Female_data_nokin,

na.action = na.omit

)

\## Aggression model without kin

Fem_model_agg_nokin <- lmer(

Cortisol ~ Age + Rank + ReproStatus + Instability + DAI + Rank*DAI + VictimRate +

(1|ChimpID) + (1|YearMonth),

data = Female_data_nokin,

na.action = na.omit

)

\# -------------------------------

\# 7. Figures for Publication

\# -------------------------------

\## 7a. Females

\# Figures showing predicted cortisol levels for female chimpanzees as a function of bond strength (DAI) and other predictors (Rank, Reproductive Status, etc.). The figure includes 95% confidence shading. See Figure 1.

figpub_females <-

plot_model(Females, type = 'pred', terms = c('DAI', 'Rank'),

title = "", axis.title = c("Female Bond Strength (DAI)", "Cortisol"),

ci.lvl = .95, legend.title = "Rank") +

theme(

panel.background = element_rect(fill = "white", color = NA),

plot.background = element_rect(fill = "white", color = NA),

panel.grid.major.x = element_blank(),

panel.grid.major.y = element_line(color = "grey90"),

panel.grid.minor = element_blank(),

axis.text = element_text(size = 11),

axis.title = element_text(size = 13)

) +

scale_color_manual(

values = c("red", "seagreen", "blue"),

labels = c("Low", "Med", "High")

) +

scale_x_continuous(expand = expansion(mult = c(0, 0)))

\# Save figure to control final resolution and aspect ratio

ggsave("figpub_females.png", figpub_females, width = 7, height = 4.2, dpi = 300)

\## 7b. Males

\# Figures showing predicted cortisol levels for male chimpanzees as a function of bond strength (DSI) and contextual variables: Instability, Mating Effort, and Aggression Rate. Each plot includes 95% confidence shading, uniform y-axis, and consistent legends. See Figure 2.

\# 7b.i. Common theme and legend settings

legend_text_size <- 10

legend_title_size <- 12

legend_key_height <- unit(0.4, "cm")

legend_box_width <- unit(1.8, "cm") # width of boxes to roughly match title length

common_theme <- theme(

panel.background = element_rect(fill = "white", color = NA),

plot.background = element_rect(fill = "white", color = NA),

panel.grid.major.x = element_blank(),

panel.grid.major.y = element_line(color = "grey90"),

panel.grid.minor = element_blank(),

axis.text = element_text(size = 11),

axis.title = element_text(size = 13),

legend.key.height = unit(0.4, "cm"),

legend.key.width = unit(1.8, "cm"), # width of colored boxes

legend.text = element_text(size = 10),

legend.title = element_text(size = 12, face = "plain"),

legend.spacing.x = unit(0.5, "cm"), # space between box and label

plot.margin = unit(c(0.5, 0.5, 0.5, 0.5), "cm")

)

\# Uniform y limits

y_limits <- c(-0.5, 0.5)

\# Helper function for two-level legends

standard_legend <- function(values, labels) {

scale_color_manual(

values = values,

labels = labels,

guide = guide_legend(

keywidth = legend_box_width

)

)

}

\# 7b.ii. Male Figures

\## (a) DSI and Instability

figpub_malesA <- plot_model(Males, type = 'pred', terms = c('DSI', 'Instability'),

title = "", axis.title = c("Bond Strength (DSI)", "Cortisol"),

ci.lvl = .95, legend.title = "Instability") +

common_theme +

standard_legend(values = c("red", "blue"), labels = c("Low", "High")) +

scale_x_continuous(expand = expansion(mult = c(0, 0))) +

coord_cartesian(ylim = y_limits)

\## (b) DSI and Mating Effort

figpub_malesB <- plot_model(Males, type = 'pred', terms = c('DSI', 'MatingEffort'),

title = "", axis.title = c("Bond Strength (DSI)", "Cortisol"),

ci.lvl = .95, legend.title = "Mating Effort") +

common_theme +

standard_legend(values = c("red", "blue"), labels = c("Low", "High")) +

scale_x_continuous(expand = expansion(mult = c(0, 0))) +

coord_cartesian(ylim = y_limits)

\## (c) DSI and Aggression Rate (3-level legend)

figpub_malesC <- plot_model(Males_Agg, type = 'pred', terms = c('DSI', 'AggressionRate'),

title = "", axis.title = c("Bond Strength (DSI)", "Cortisol"),

ci.lvl = .95, legend.title = "Aggression Rate") +

common_theme +

scale_color_manual(

values = c("red", "seagreen", "blue"),

labels = c("Low", "Med", "High"),

guide = guide_legend(

keywidth = legend_box_width

)

) +

scale_x_continuous(expand = expansion(mult = c(0, 0))) +

coord_cartesian(ylim = y_limits)

\# 7b.iii. Add figure labels

figpub_malesA <- ggdraw(figpub_malesA) + draw_label("(a)", x = 0, y = 1, hjust = -0.1, vjust = 1.1, fontface = "bold", size = 12)

figpub_malesB <- ggdraw(figpub_malesB) + draw_label("(b)", x = 0, y = 1, hjust = -0.1, vjust = 1.1, fontface = "bold", size = 12)

figpub_malesC <- ggdraw(figpub_malesC) + draw_label("(c)", x = 0, y = 1, hjust = -0.1, vjust = 1.1, fontface = "bold", size = 12)

\# 7b.iv. Combine all male figures vertically

figpub_males <- plot_grid(

figpub_malesA,

figpub_malesB,

figpub_malesC,

ncol = 1,

align = "v",

rel_heights = c(2, 2, 2)

)

\# Save figure to control final resolution and aspect ratio

ggsave("figpubs_males.png", figpub_males, width = 6, height = 9, dpi = 300, bg = "white")

\# -------------------------------

\# 8. Session Info

\# -------------------------------

\# Print session info for reproducibility

sessionInfo()

```
